# Cardiac and gastric interoception have distinct neural substrates

**DOI:** 10.1101/2022.02.18.480981

**Authors:** Yusuke Haruki, Kenji Ogawa

**Affiliations:** Department of Psychology, Graduate School of Humanities and Human Sciences, Hokkaido University, Kita 10, Nishi 7, Kita-ku, Sapporo 060-0810, Japan

## Abstract

Interoception, or an awareness of the internal body state, guides agents in adaptive behavior by informing them of ongoing bodily signals, such as heart rate or energy status. However, it is still unclear whether the human brain represents the differences in the subjective experience of interoception differently. Hence, we directly compared the neural activation for cardiac (awareness related to heartbeats) and gastric (awareness related to the stomach) interoception in the same population (healthy human, N = 31). Participants were asked to focus on their heart and stomach sensations to be aware of interoception in a magnetic resonance imaging scanner. The results indicated that neural activation underlying gastric interoception encompassed larger brain regions, including the occipitotemporal visual cortices, bilateral primary motor cortex, primary somatosensory cortex, left orbitofrontal cortex, and bilateral hippocampal regions. Cardiac interoception, however, selectively activated the right anterior insula extending to the frontal operculum more compared to gastric interoception. Moreover, our detailed analyses focusing on the insula, the most relevant region for interoception, revealed that the left dorsal middle insula encoded cardiac and gastric interoception in different activation patterns but not the posterior insula. Our results demonstrate that cardiac and gastric interoception have distinct neural substrates; in particular, the selective brain activation may reflect differences in the functional roles of cardiac and gastric interoception.

**Significance statement:** Interoception, subjective senses that arise from within the body, plays a critical role in maintaining adaptive behavior by informing of the ongoing bodily states, such as heart rate and energy status. Although interoception has various characteristics depending on its source signals, previous neuroimaging studies have extensively used cardiac interoception (senses related to heartbeats), making it unclear whether the brain differently encodes diverse experiences of interoception. Here, we demonstrate that cardiac interoception and gastric interoception (senses related to the stomach) have distinct neural substrates by combining mass-univariate analysis with multivoxel pattern analysis for fMRI data. Our findings suggest that the selective brain activation may reflect differences in the functional roles of cardiac and gastric interoception.

## Introduction

Subjective experiences of one’s internal bodily state are referred to as interoception or interoceptive awareness (Brewer et al., 2021; Khalsa et al., 2018). Such senses indicate ongoing physiological changes, guiding agents in adaptive behavior (Paulus et al., 2019; Quigley et al., 2021). In this paper, we use the term interoception to describe the phenomenal experience of bodily states. Examples of interoception changing the way agents sample sensory information and act in the external world are as follows: thirst compels agents to prioritize finding and drinking water, heartbeat perception signals physiological arousal and increases vigilance, and an empty stomach informs when and where to eat. Although interoception serves multiple functional roles depending on phenomenal differences, it remains unclear how the human brain represents diverse experiences of interoception (Azzalini et al., 2019). We thus used functional magnetic resonance imaging (fMRI) to focus on the brain activation of cardiac interoception (i.e., awareness of one’s heartbeat sensation) and gastric interoception (awareness of the stomach sensation) that have been suggested to serve different functional roles.

In situations where people become aware of their heartbeats, the heartbeat signals and cardiac interoception inform of changes in physiological arousal (Paulus et al., 2019). For example, accelerated false cardiac feedback (i.e., exaggerated cardiac interoception) has been found to alter the emotional salience of neutral faces (Gray et al., 2007) and perceived physical effort (Iodice et al., 2019) by enhancing physiological arousal. Additionally, gastric interoception such as fullness or hunger modulates foraging and feeding behavior (Wang et al., 2008). In fact, people with an eating disorder often show an altered gastric interoception (Van Dyck et al., 2021; Walsh et al., 2003), which may cause their abnormal eating behavior. Despite considerable differences in their functional roles, extant research has failed to establish whether the brain represents cardiac and gastric interoception differently.

Invasiveness to manipulate interoception has limited the detailed functional brain mapping of interoception. The brain regions responding to changes in visceral signals and interoception have been studied using a physical stimulation on viscera (Jarrahi et al., 2015; Lu et al., 2004) or a pharmacological disturbance (Hassanpour et al., 2018, 2016) during fMRI scanning. However, such techniques cannot be combined in a single study because of their invasiveness, making it difficult to compare the brain activation for diverse interoception. Another way to map interoception is to use an interoceptive attention paradigm—asking participants to direct their attention toward sensations originating from within the body. Importantly, although exteroceptive information such as cutaneous sensations may contribute to interoception (Khalsa et al., 2009), interoceptive attention has been found to elicit a robust activation of visceral sensory areas, the insula (Haruki and Ogawa, 2021; Klabunde et al., 2019; Wiebking et al., 2014). Particularly, researchers have consistently associated the activation in the right anterior insula with interoceptive attention (Simmons et al., 2013; Zaki et al., 2012) and objective accuracy of cardiac interoception (Caseras et al., 2013; Critchley et al., 2004; Pollatos et al., 2007). These results appear to support an influential neuroanatomical model of interoception where the posterior insula receives an initial cortical input of visceral signals while the right anterior insula represents an awareness of internal bodily states (Craig, 2009, 2011; Evrard, 2019).

Previous studies using the interoceptive attention paradigm suffer from the limitation of having extensively used heartbeat attention (i.e., cardiac interoception) over other modalities of interoception. Even when the attentional focus on gastric interoception was deployed, direct comparison within the modality of interoception has not been discussed (DeVille et al., 2020; Kerr et al., 2016; Simmons et al., 2013). Therefore, it is still unclear whether neural encodings of the various interoception differ, limiting the theoretical advances in this field. To address this issue, we directly compared the neural activation for cardiac and gastric interoception using the interoceptive attention paradigm in a healthy population. We hypothesized that cardiac and gastric interoception would elicit distinct neural activations, which would be modulated depending on their functional role. That is, cardiac interoception might activate the regions underlying physiological arousal; brain areas that modulate feeding and foraging behavior might show enhanced activation in gastric interoception. Moreover, we considered that the insula, the most relevant region for interoception (Craig, 2009; Critchley and Harrison, 2013), would show a subregion-specific representation of cardiac and gastric interoception. To test this idea, we performed a region of interest (ROI) analysis focusing on the insula by combining multivoxel pattern analysis (MVPA) with a basic comparison of neural activation.

## Materials and methods

### Participants

A total of 35 right-handed people participated in the experiment (15 women). One participant (woman) requested an interruption during the experiment, yielding incomplete data. Of the remaining participants, three (one man and two women) were excluded from the analysis because their maximum head movement was more than 3 mm during the experiment. Thus, the final analysis included 31 participants (12 women) who were 21.61 ± 2.45 years old (range: 20 to 31). Their handedness was assessed by a modified version of the Edinburgh Handedness Inventory for Japanese participants (Hatta and Nakatsuka, 1975). The sample size deemed sufficient for determining reliable brain activation was calculated based on previous studies that have used a similar task for interoception (Haruki and Ogawa, 2021; Wiebking and Northoff, 2015). Written informed consent was obtained from all participants. This study was carried out in accordance with the Declaration of Helsinki and all its future amendments. The Ethics Committee of Hokkaido University approved the experimental protocol.

### Task procedures

To quantify the brain activation for cardiac and gastric interoception noninvasively, we asked participants to perform an established interoceptive attention task in an MRI scanner. We used the task procedure that elicited robust brain activation for interoception (DeVille et al., 2020; Kerr et al., 2016; Simmons et al., 2013), which was modified to suit our purpose and to increase the statistical power. The present task was a simple block design that had three conditions with resting periods: heart attention (for cardiac interoception), stomach attention (for gastric interoception), and visual attention (for control). Participants were asked to focus on sensations from their heart, stomach, or visual stimuli to be aware of the slight sensations for each condition (Figure 1). In the heart attention and stomach attention trial, the words “HEART” and “STOMACH” were presented for 10 s each on the screen to allow participants to realize which sensation they were focusing on. In the visual attention trial, the word “TARGET” was presented on the screen for 10 s, with the color of the word gradually and slightly fading from black to gray. The color changed every 1.5 s for a total of five times. Therefore, in each task trial, participants directed their attention to a particular, vague sensation without any salient stimulus. During the rest period, participants watched a fixation crossbar for 12 s with their eyes kept open. Preparing for the next task trial was prohibited.

**Figure 1.**
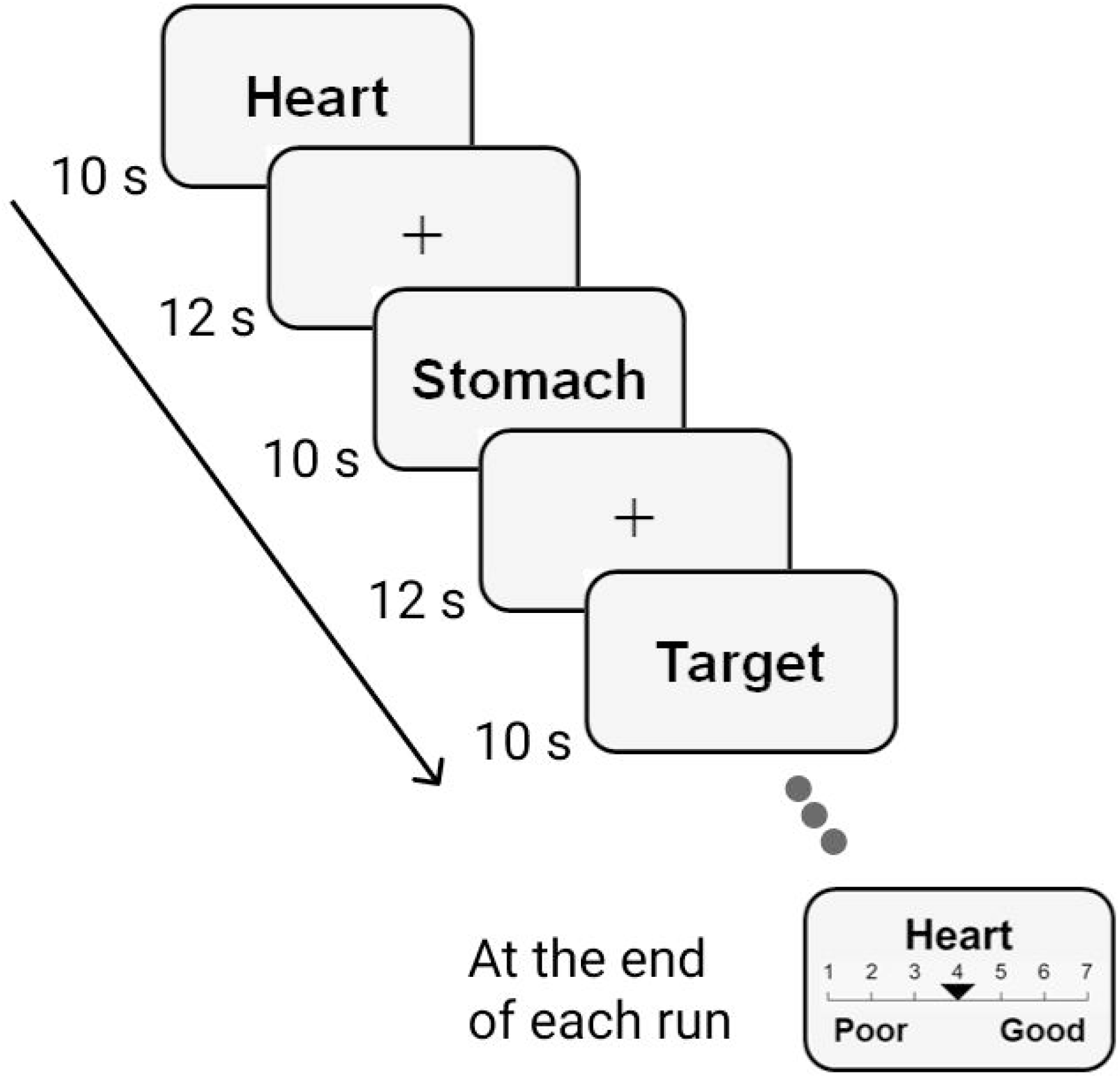
Schematical presentation of the task procedure. An example of the time course of the functional magnetic resonance imaging (fMRI) task is depicted. Our block design fMRI task included three types of task trials (cardiac interoception, gastric interoception, and visual) that lasted 10 s each, followed by rest (12 s). Participants were asked to focus on sensations arising from the heart, stomach, and color change of the word, according to the type of task. At the end of each run, participants rated the subjective intensity of the sensations for each condition using a Likert scale ranging from 1 (not intense at all) to 7 (extremely intense).

Each condition of the task trial was presented five times per run: a single run included a total of 15 task trials and 15 rest periods. The order of task trials was pseudo-randomized, but the same trial was not presented three times in succession. At the end of each run, participants rated the subjective intensity of the sensations (heart, stomach, and visual) throughout the run using a Likert scale ranging from 1 (not intense at all) to 7 (extremely intense). An approximately 5.5 min run was repeated five times; thus, we obtained 25 data points (125 volumes) for each condition per participant. This procedure was designed to include larger number of trials compared with previous studies (DeVille et al., 2020; Kerr et al., 2016) because we needed sufficient number of task trials to perform MVPA. Before the first run began, all participants underwent a resting-state scanning that lasted for 5 min, which was not analyzed in the present study. Stimuli were presented on a liquid crystal display and projected onto a custom-made viewing screen. Participants took a supine position in the scanner and viewed the screen via a mirror. Participants experienced a practice trial and its intensity report (for each condition) to learn how to perform the task and report the subjective intensity before entering in the MRI scanner. Through the practice trials, we verbally confirmed that participants successfully directed their attention to each sensation depending on the condition.

### MRI acquisition

All scans were performed on a Siemens (Erlangen, Germany) 3-Tesla Prisma scanner with a 64-channel head coil at Hokkaido University. T2*-weighted echo planar imaging (EPI) was used to acquire a total of 168 scans per run, with a gradient echo EPI sequence. The first three scans within each session were discarded in order to allow for T1 equilibration. The scanning parameters used were as follows: repetition time (TR), 2,000 ms; echo time (TE), 30 ms; flip angle (FA), 90°; field of view (FOV), 192 × 192 mm; matrix, 94 × 94; 32 axial slices; and slice thickness, 3.500 mm, with a 0.875 mm gap. Thus, the voxel size was 2.042 × 2.042 × 4.375 mm. T1-weighted anatomical imaging with an MP-RAGE sequence was performed using the following parameters: TR, 2,300 ms; TE, 2.32 ms; FA, 8°; FOV, 256 × 256 mm; matrix, 256 × 256; 192 axial slices; and slice thickness, 1 mm without a gap.

### Preprocessing of fMRI data

All image preprocessing was performed using the SPM12 software (Wellcome Department of Cognitive Neurology, http://www.fil.ion.ucl.ac.uk/spm). All functional images were initially realigned to adjust for motion-related artifacts. Volume-based realignment was performed by co-registering images using rigid-body transformation to minimize the squared differences between volumes. The realigned images were then spatially normalized with the Montreal Neurological Institute template based on the affine and non-linear registration of co-registered T1-weighted anatomical images. They were resampled into 3-mm-cube voxels with sinc interpolation. Images were spatially smoothed using a Gaussian kernel of 6×6×6-mm full width at half-maximum. The images used for MVPA were not smoothed to avoid blurring the information contained in the multi-voxel activity pattern.

## Statistical analysis

### Subjective intensity of the sensations

We used R (version 4.0.3) for all our statistical inference except functional imaging data. First, we tested whether the subjective intensity of sensations differed between modalities (cardiac, gastric, and visual). The linear mixed-effects (LME) model analysis implemented in the lme4 package (Bates et al., 2015) was performed. With all the data obtained (N = 465, 3 trial types for 5 runs for 31 participants) as a dependent variable, we modeled the type of sensation as the fixed effect. Random intercept and slope for the effects of participants and random intercept for the runs were modeled as random effects, ensuring the maximal random structure for our models (Barr et al., 2013). The LME enabled us to avoid averaging the values across five runs compared to the traditional analysis of variance (ANOVA) (Baayen et al., 2008). The parameter was estimated using the restricted maximum likelihood method, with the degrees of freedom estimated using the Satterthwaite method.

### Whole-brain activation

We first evaluated the brain regions activated in each condition (heart attention, stomach attention, and visual attention) using a generalized linear model (GLM). Individual-level GLM included three regressors of interest for each condition as a separate box-car function that was convolved with the canonical hemodynamic response function. The rest period was used as a baseline. To reduce motion-related artifacts, six motion parameters were included as nuisance regressors. By combining the three conditions, we obtained six contrast images (heart attention compared to stomach attention, heart compared to visual, stomach compared to visual, and their opposite contrasts) for each participant. We then performed a group-level random effects analysis for these images using a one-sample t-test. Through these statistical inferences, we directly compared the brain activation for each condition across the whole brain.

Moreover, we performed a group-level analysis of covariance (ANCOVA) to exclude the effect of the subjective ratings of stimulus intensity on brain activation. This was because we considered that the differences in the subjective intensity may affect the brain activation pattern, independently of the object of attentional focus. Using the image of interoception contrasted to the visual control, we modeled individual subjective ratings for each condition, averaged for five runs, as covariates. Then, ANCOVA excluding the effects of subjective rating as nuisance was performed. By doing so, we assessed the brain activation of cardiac interoception contrasted to visual attention without the effect of individual differences in subjective heartbeat intensity and gastric interoception contrasted to visual without the subjective intensity of stomach sensation. Furthermore, we explored brain activations varying as a function of the subjective intensity; a regression analysis with the subjective intensity as a covariate of interest was performed using the same model used in ANCOVA. The voxel-level threshold was set to *p* < .001 (uncorrected), and the cluster-level threshold was set to *p* < .05, corrected for family-wise error (FWE).

### ROI analysis of the subregions of the insula

We tested whether the subdivisions of the insula, the primary interoceptive cortex that receives the initial cortical input of visceral signals (Craig, 2002), had a different representation for cardiac and gastric interoception at the voxel level. In particular, we first performed MVPA that allows us to investigate more sophisticated neural representation compared with conventional analysis (Norman et al., 2006), suspecting that conventional analysis might fail to detect the activation difference in the insula. The ROIs were defined as the anatomical subdivisions of the insula in Hammersmith brain atlases (Brain Development, www.brain-development.org). These images were constructed as a 3D probabilistic atlas using in vivo T1 MR images, including anatomical structures commonly seen in the human insula: the anterior short gyrus (ASG; the most dorsal anterior portion of the insula), middle short gyrus (MSG; the dorsal mid-anterior), posterior short gyrus (PSG; the dorsal mid-posterior), anterior inferior cortex (AIC; the ventral anterior), anterior long gyrus (ALG; the dorsal posterior), and posterior long gyrus (PLG; the ventral posterior) (Faillenot et al., 2017). Importantly, these ROIs have been confirmed to show a subregion-specific activation pattern for cardiac interoception (Haruki and Ogawa, 2021).

MVPA for cardiac and gastric interoception was performed with a two-class classifier based on a linear support vector machine (SVM) implemented in LIBSVM (http://www.csie.ntu.edu.tw/~cjlin/libsvm/). We first created another individual-level GLM apart from the whole-brain analyses that included 15 task trials as independent regressors per run with six motion parameters as nuisance regressors. By doing so, we obtained parameter estimates of all voxels in each ROI for a total of 75 trials per participant, labeling them as heart, stomach, or visual, depending on the condition of each trial. We then trained SVM to classify the identity of the brain activation pattern of cardiac and gastric interoception using these parameter estimates as inputs to the SVM. Individual-level classification accuracy was estimated with a five-fold “leave-one-out” cross-validation to avoid overfitting. This procedure uses inputs in four runs as training data and inputs in one remaining run as test data to be classified, which was repeated five times for all possible combinations. The averaged classification accuracy across five repetitions of the tests was computed for each ROI for each participant independently. We used a default hyperparameter (a fixed regularization parameter C = 1). Parameter estimates of the trial were not used as input to the classifier. One-sample t-tests were performed to test whether the activation patterns for cardiac and gastric interoception in each subregion were classifiable using SVM. That is, we tested whether the computed classification accuracy was higher than the chance level (50%) at the group level, separately for each ROI. Because we used 12 ROIs (six anatomical regions for both hemispheres), the calculated *p* values were corrected for Benjamini and Hockberg’s false discovery rate (FDR; Benjamini & Hochberg, 1995).

We also compared the difference in the averaged activation in the ROIs among each condition because we suspected that the classification accuracy calculated by MVPA could merely reflect the mean signal change in the ROIs. First, the mean signal changes in each ROI were extracted for each participant, separately for the cardiac and gastric attention condition. Then, we performed a repeated-measures ANOVA on these values with the condition (cardiac and gastric) and anatomical location (the ASG, MSG, PSG, AIC, ALG, and PLG) as within-factors, independently for each hemisphere. Multiple comparison correction for post-hoc analyses was performed using Shaffer’s modified sequentially rejective Bonferroni procedure implemented in R. The combined use of MVPA and direct comparisons of the averaged signal change allowed detailed investigation of how the human insula represents interoception specific to the cardiac and gastric domain.

Furthermore, we tested whether averaged activation for interoception and exteroception differed in the insula using a similar procedure in the case of cardiac and gastric interoception. We extracted mean signal changes for interoception, the averaged activation across heart and stomach attention, and exteroception (visual attention) in each ROI. Then, repeated-measures ANOVA with the condition (interoception and exteroception) and the anatomical location as within-factors was performed separately for each hemisphere.

## Results

### Subjective intensity of the sensations

We performed an LME model analysis for the subjective rating of the intensity of each sensation. The LME included the type of stimulus (heart, stomach, and visual) as the main factor; random intercept and slope for the effects of the participant and random slope for the sequence of runs were modeled as random effects. The subjective intensity of visual stimuli was rated the highest (marginal mean contrasted to heart = 0.90, *t*(30.00) = 4.51, *p* < .001) and that of the stomach the lowest (marginal mean = -0.41, *t*(30.00) = -2.22, *p* = .03) among the three stimuli (Table 1, Figure 3B).

**Table 1.**
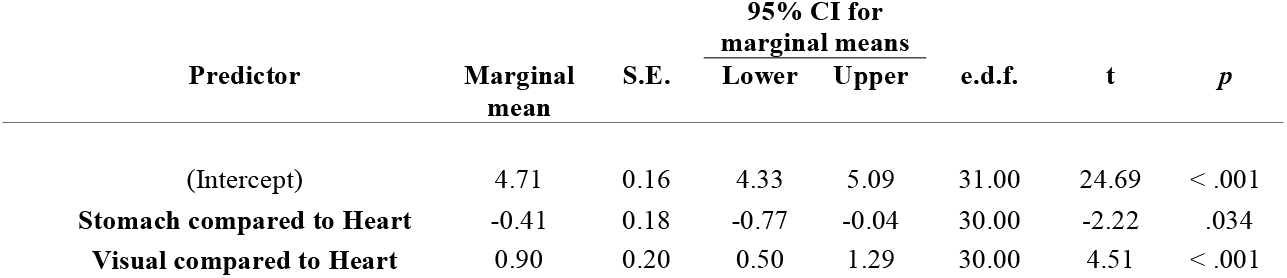
Subjective intensity of sensations in each condition. The linear mixed-effects model included the condition as the fixed effect. The random structure was maximal for appropriate inference: random intercept and slope for the effect of subject and random intercept for the effect of run sequence were modeled as random effects. S.E., standard error for marginal means; CI, confidence interval; e.d.f., estimated degrees of freedom

| Predictor | Marginal mean | S.E. | 95% CI for marginal means |  | e.d.f. | t | p |
| --- | --- | --- | --- | --- | --- | --- | --- |
|  |  |  | Lower | Upper |  |  |  |
| (Intercept) | 4.71 | 0.16 | 4.33 | 5.09 | 31.00 | 24.69 | < .001 |
| **Stomach compared to Heart** | -0.41 | 0.18 | -0.77 | -0.04 | 30.00 | -2.22 | .034 |
| **Visual compared to Heart** | 0.90 | 0.20 | 0.50 | 1.29 | 30.00 | 4.51 | < .001 |

### Whole-brain activation for cardiac and gastric interoception

We directly compared whole-brain activation across three conditions (cardiac interoception, gastric interoception, and control vision). We found that gastric interoception activated larger brain areas, including the occipitotemporal visual cortices, bilateral primary motor, primary somatosensory, left orbitofrontal, and bilateral posterior hippocampus, compared to cardiac interoception. In contrast, the right dorsal anterior insula extending to the frontal operculum only showed higher activation in cardiac than gastric interoception (Figure 2A, Table 2). In particular, the medial visual area that was enhanced during gastric interoception largely overlapped with the “gastric network” that showed a temporal coupling with the gastric basal rhythm (Figure 2B) (Rebollo and Tallon-Baudry, 2022). Moreover, by comparing cardiac and gastric interoception to the visual control condition, we found that cardiac and gastric interoception activated similar brain regions, including the insula, frontal operculum, parietal operculum, middle cingulate, and supplementary motor area; the results were highly comparable to previous studies (Haruki and Ogawa, 2021; Simmons et al., 2013; Tan et al., 2018). A notable exception was that the hippocampus and medial visual areas were activated only in gastric interoception (Figure 3A). The visual control condition contrasted to cardiac and gastric interoception activated brain regions critical for visual attention, such as the middle frontal gyrus, superior parietal lobule, posterior thalamus (geniculate nucleus), and lateral visual association area (Dosenbach et al., 2007; Mayer et al., 2007), supporting the validity of our experimental design (Figure 3A). Furthermore, we confirmed that the brain activation elicited by cardiac and gastric interoception was almost unaffected by differences in subjective stimulus intensity by performing a multiple regression analysis excluding the effects of subjective ratings of stimulus intensity. The results of the multiple regression were fairly comparable to the brain activation obtained in cardiac and gastric interoception contrasted to visual control (Figure 3C). There was no brain activation covarying with the subjective intensity rating even at a more moderate threshold (cluster-level *p* < .05, uncorrected). All results are reported with a voxel-level threshold of *p* < .001 (uncorrected) with cluster size correction for *p* values < .05 (FWE).

**Table 2.**
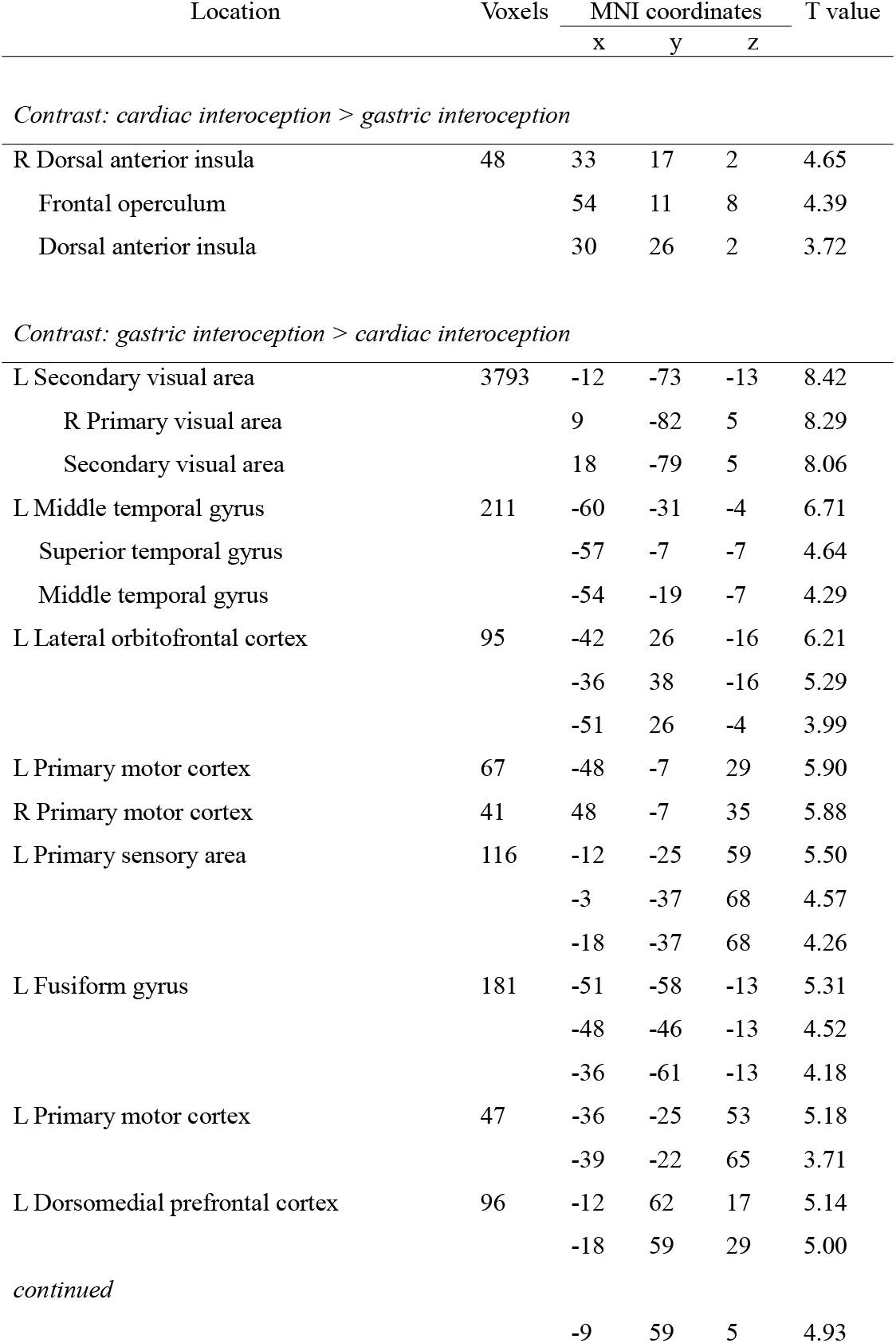
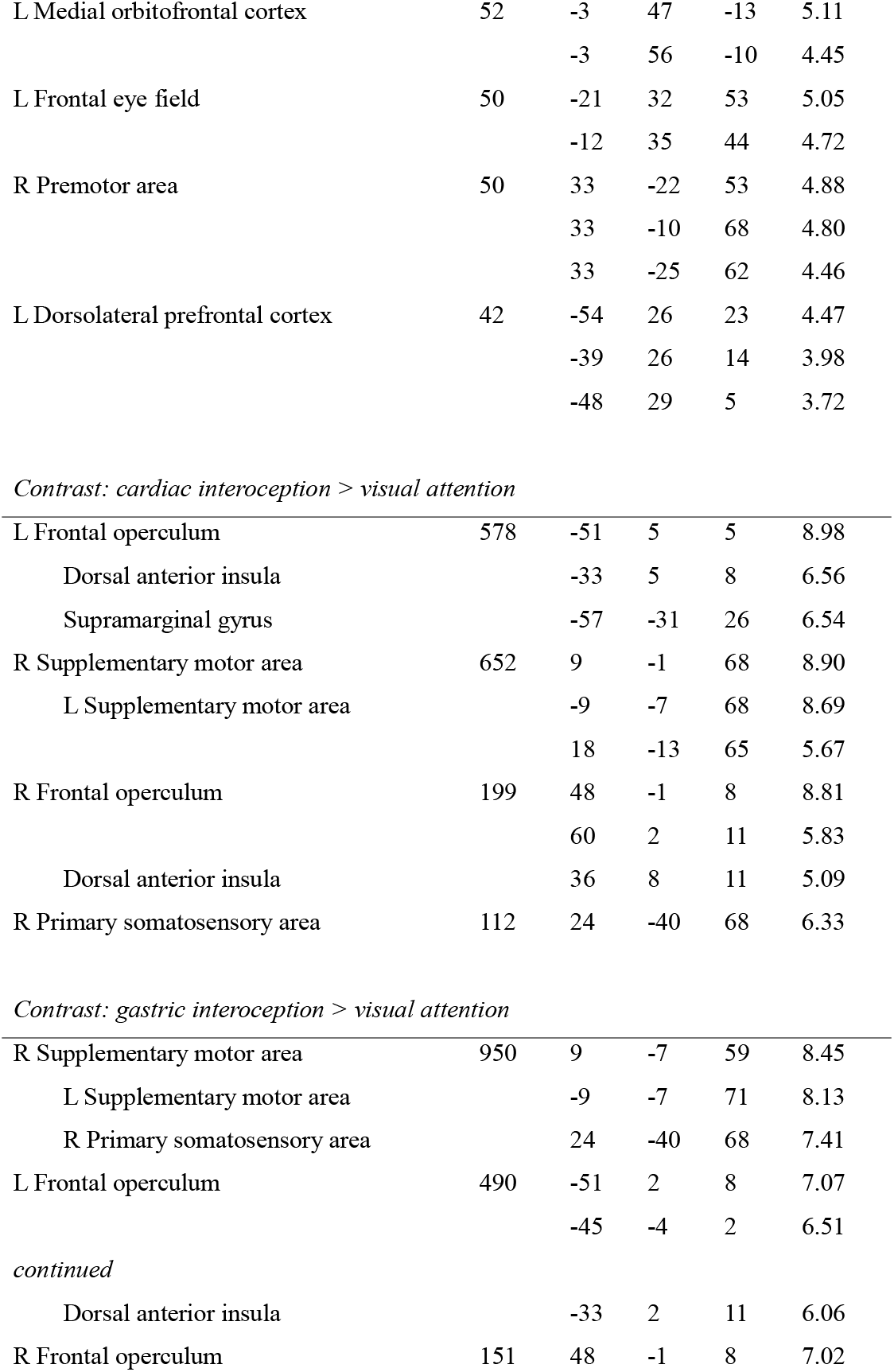
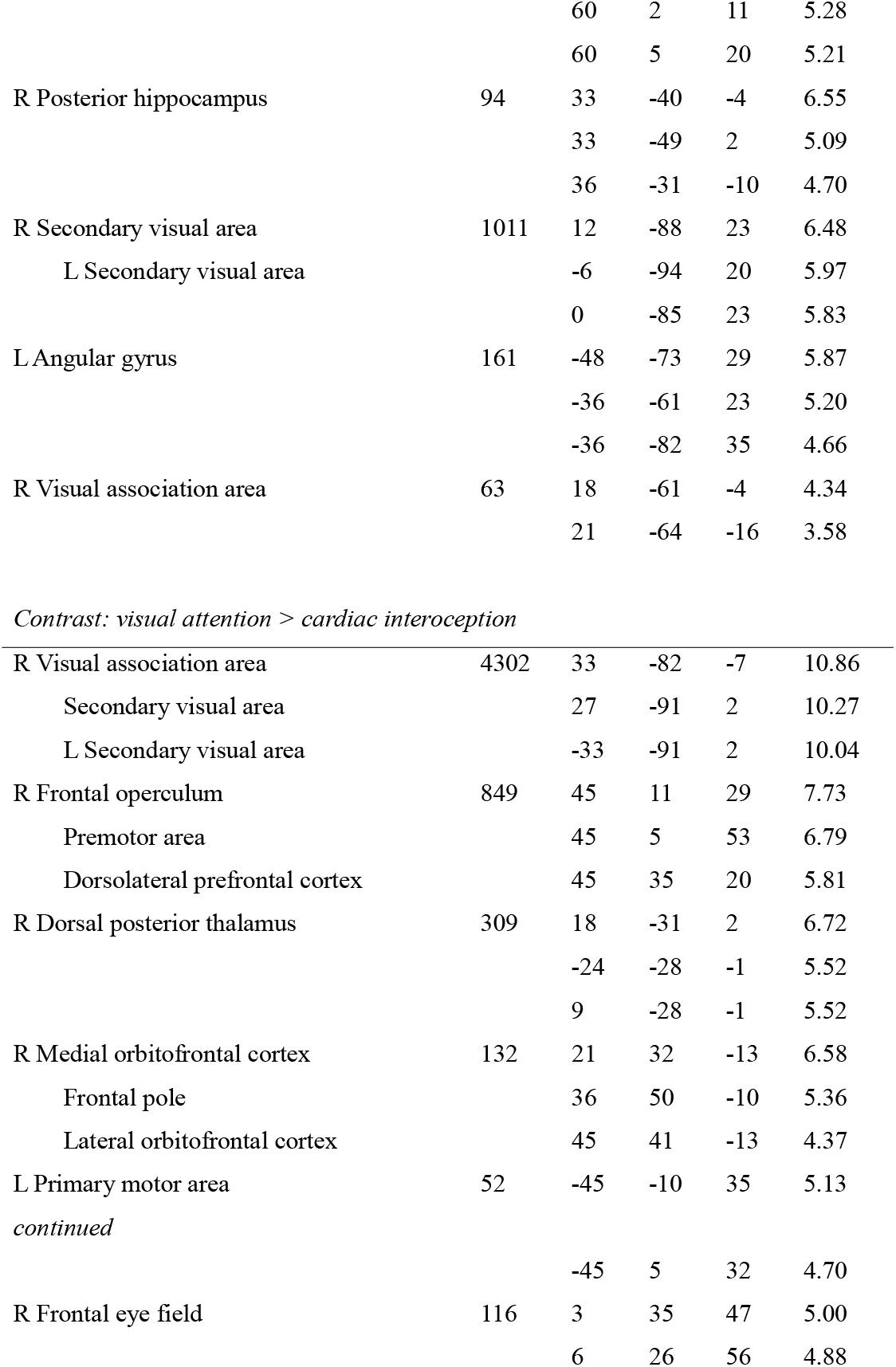
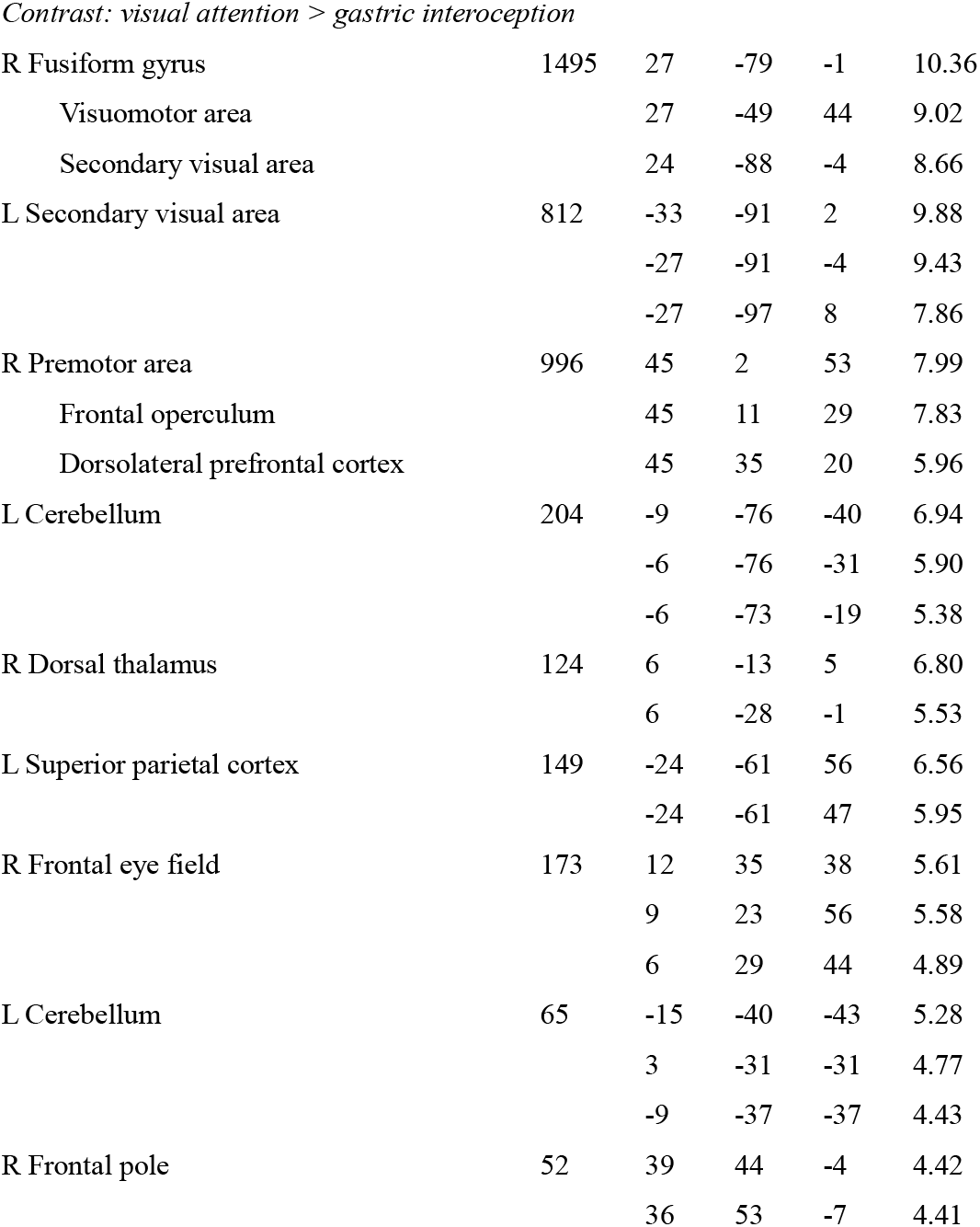
Anatomical regions, peak voxel coordinates, and t-values of observed activations. The peak-level threshold was set to *p* < .001 and the cluster size was also corrected for extent threshold of *p* < .05 (corrected for family-wise error). L, left hemisphere; MNI, Montreal Neurological Institute; R, right hemisphere

**Figure 2.**
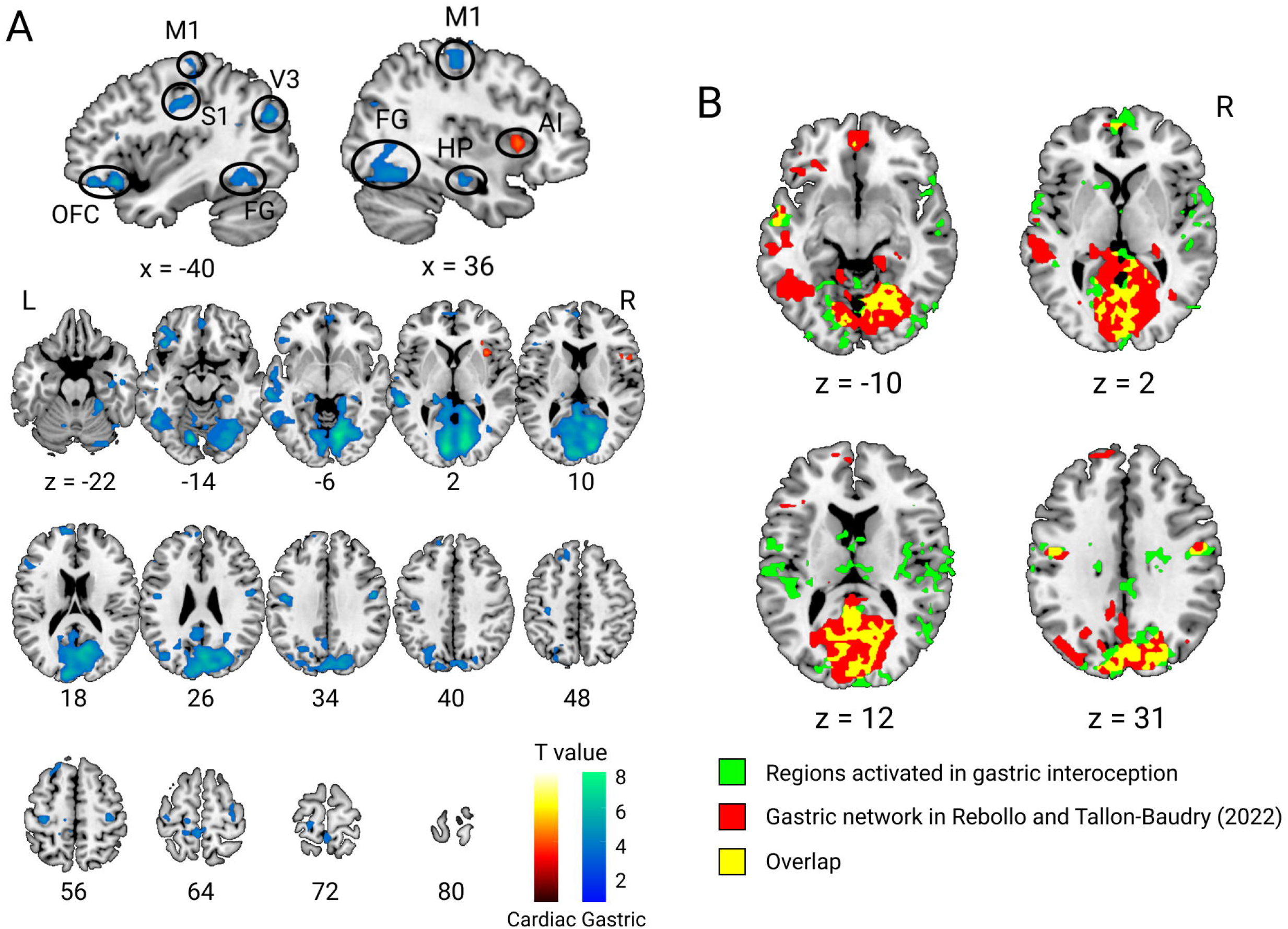
Whole-brain activation for cardiac and gastric interoception. A). Significant activation in cardiac interoception contrasted with gastric interoception (warm color) and vice versa (cold color) are depicted with a height threshold of *p* < .001 (uncorrected) and an extent threshold of *p* < .05 (corrected for family-wise error, corresponding to more than 42 voxels). Sagittal views show that cardiac and gastric interoception activated the brain region relevant to each interoception. Sensorimotor regions (S1: primary sensory, M1: primary motor, V3: visual area 3, FG: fusiform gyrus), orbitofrontal cortex (OFC), and hippocampus (HP) showed enhanced activation in gastric interoception, while the right anterior insula (AI) was activated in cardiac interoception. B). The internally extended visual area that was activated in gastric interoception largely overlapped with the brain regions that have been shown to couple with the gastric basal rhythm (Rebollo and Tallon-Baudry, 2022). L, left hemisphere; R, right hemisphere. All figures are shown in axial slices with z denoting locations in the Montreal Neurological Institute coordinates.

**Figure 3.**
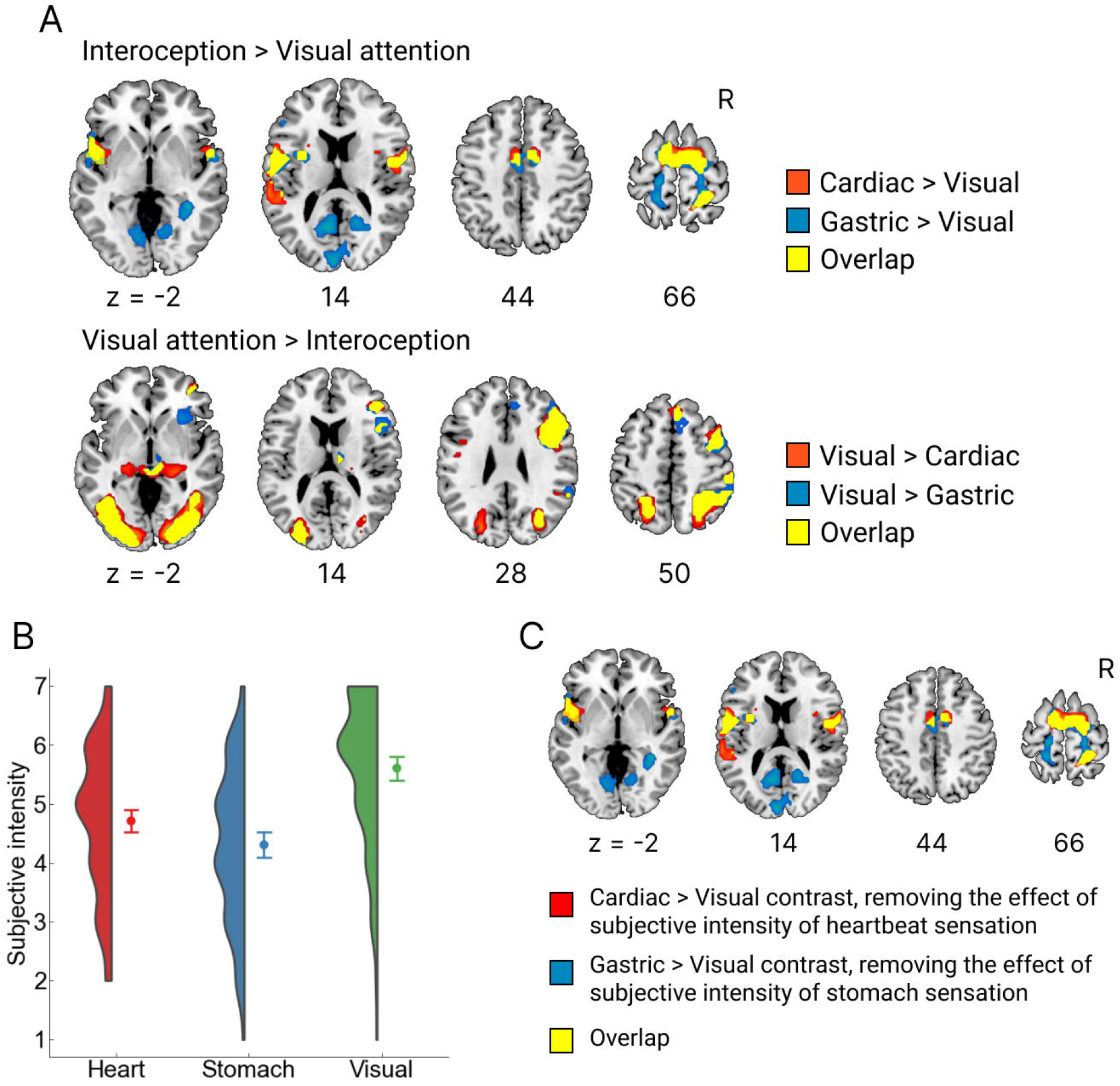
Brain activation for cardiac and gastric interoception compared with control. A). When compared with the visual attention condition, cardiac and gastric interoception showed similar activation patterns, such as activations in the insula and supplemental motor area and deactivations in the right middle frontal, superior parietal, posterior thalamus, and lateral visual association area. However, activations in the medial visual cortex and hippocampus were found only in gastric interoception compared with visual attention. Height threshold was set to *p* < .001 (uncorrected) with an extent threshold of *p* < .05 (corrected for family-wise error), corresponding to more than 52 voxels. B). Distribution and mean scores of the subjective report of stimulus intensity are plotted separately for each condition. Participants reported the intensity at the end of scanning run for each condition, yielding 155 data (31 participants for 5 runs) for each condition. Our maximal linear mixed-effects model analysis revealed that the visual stimulus (slight change of word color) was rated the most intense (regression coefficient compared to heart = 0.90, t_30.00_ = 4.51, *p* < .001); the stomach sensation was rated the least intense (regression coefficient = -0.41, t_30.00_ = -2.22, *p* = .034). Error bars show 95% confidence interval while the half-violin plot represents the kernel density estimation. C). The brain activation in cardiac and gastric interoception was not varied as a function of the subjective report of stimulus intensity. We performed a multiple regression analysis that excluded the effect of subjective reports from brain activation, revealing almost the same activation patterns compared with activations reported in A). Height threshold was set to *p* < .001 (uncorrected) with an extent threshold of *p* < .05 (corrected for family-wise error), corresponding to more than 52 voxels. R, right hemisphere. All figures are shown in axial slices with z denoting locations in the Montreal Neurological Institute coordinates

### Multivoxel pattern classification for cardiac and gastric interoception in the insula

We performed MVPA using an SVM classifier that distinguished the activation patterns in the insula between cardiac and gastric interoception. Activation patterns in the anatomical subregions of the insula (Faillenot et al., 2017, Figure 4A) were used as inputs to the SVM. We found that classification accuracy in the left PSG, which corresponds to the dorsal middle insula, was significantly higher than the chance level (50%) (mean classification accuracy = 56.32, t_30_ = 3.39, corrected *p*-value for FDR = .024), suggesting that the region had distinct representation for cardiac and gastric interoception. The right ASG (mean = 53.48, t_30_ = 2.56, corrected *p* = .089), left ASG (mean = 54.26, t_30_ = 2.28, corrected *p* = .089), and left MSG (mean = 53.42, t_30_ = 2.34, corrected *p* = .089) showed a marginally significant classification accuracy above the chance level; all of them would correspond to the dorsal mid-anterior insula. The other subdivisions of the insula such as posterior or ventral subdivisions showed no significant classification accuracy above the chance level (corrected *p*s > .27) (Figure 4B).

**Figure 4.**
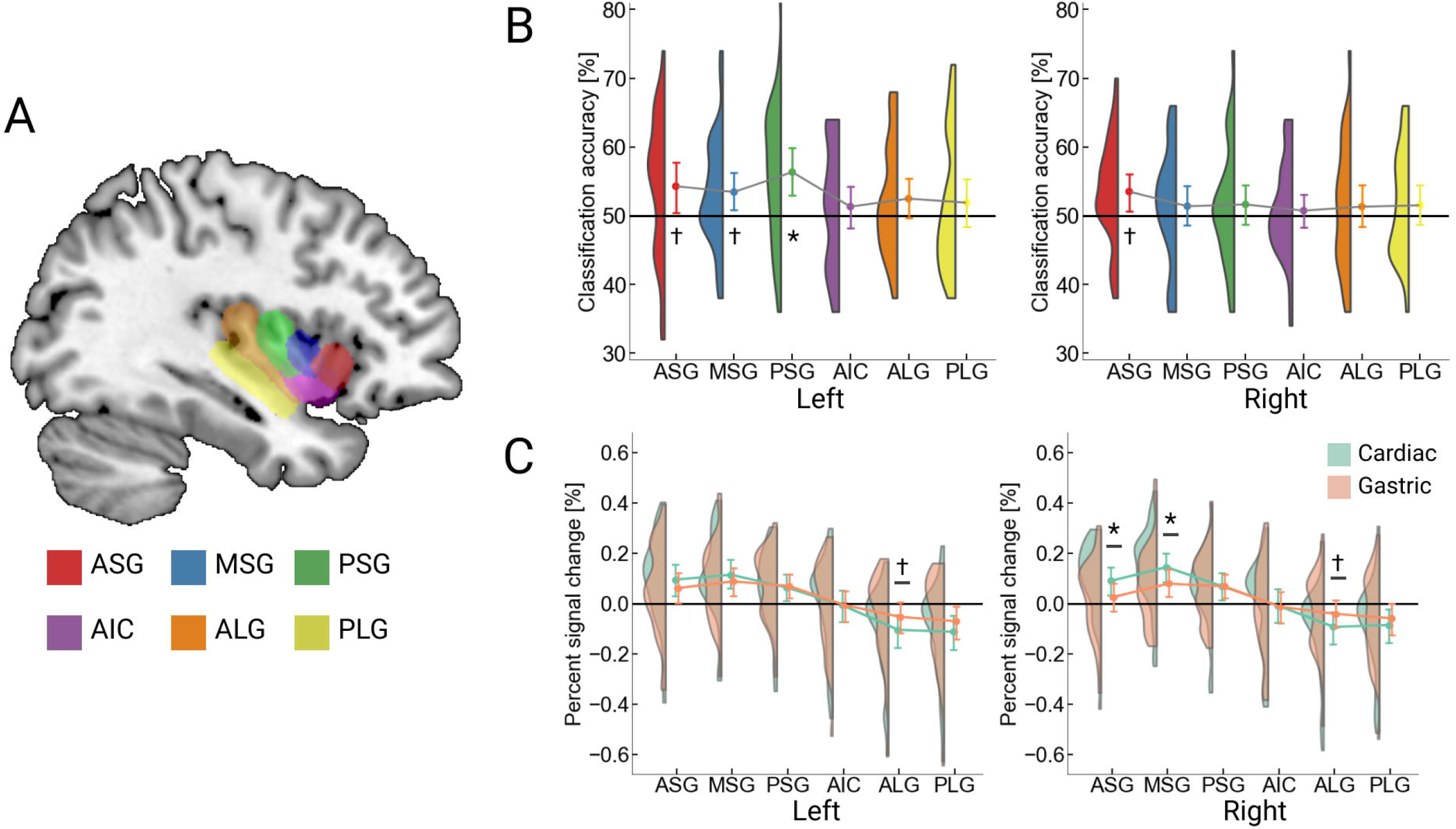
Comparison of activation for cardiac and gastric interoception in the subdivisions of insula. A). The anatomical subdivisions of the insula (Faillenot et al., 2017) are presented. We adopted six regions (the anterior short gyrus: ASG, middle short gyrus: MSG, posterior short gyrus: PSG, anterior insular cortex: AIC, anterior long gyrus: ALG, and posterior long gyrus: PLG) for both hemispheres. B). The results of multivoxel pattern analysis are depicted, with the left panel showing the results of left insula subdivision and the right panel showing the right insula. The left MSG exhibited significantly higher classification accuracy above 50% chance level. A marginally significant classification accuracy (i.e., *p* < .1) was observed in the right ASG, left ASG, and left MSG. The point plot represents the mean classification accuracy with 95% confidence interval while the half-violin plot represents the kernel density estimation. C). The results of direct comparison of the signal strength between cardiac and gastric interoception are plotted. Post-hoc analysis of repeated-measures analysis of variance revealed that the right ASG and MSG showed significantly higher activation in cardiac interoception than gastric interoception. There was a marginally significant effect that implies stronger activation in gastric than cardiac interoception in the left ALG and right ALG. The point plots represent the mean signal change with 95% confidence interval while the half-violin plot represents the kernel density estimation (green for cardiac, orange for gastric interoception). * *p* < .05; † *p* < .10 (corrected for false discovery rate). As extended data, we performed a similar analysis focusing the subdivisions of the insula for averaged brain activation in interoception and exteroception (Extended Data Figure 4-1).

**Figure 4-1.**
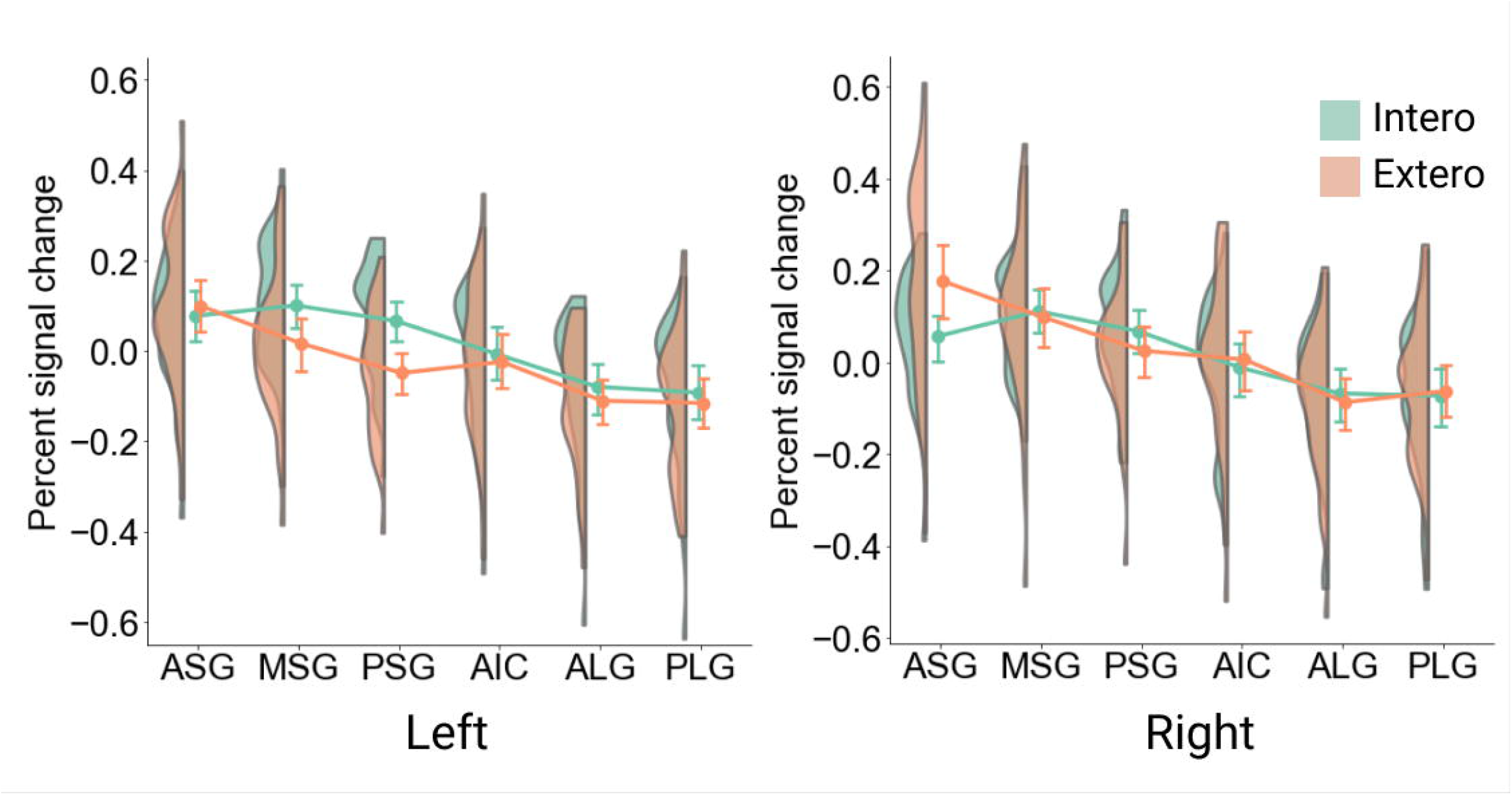
Comparison of activation for interoception and exteroception in the subdivisions of insula. The results of direct comparison of the signal strength between interoception (averaged for heartbeat attention and stomach attention) and exteroception (visual attention) are plotted. Post-hoc analysis of repeated-measures analysis of variance revealed that the right ASG showed significantly higher activation in exteroception than interoception, whereas the left MSG and PSG were more activated in interoception. The point plots represent the mean signal change with 95% confidence interval while the half-violin plots represent the kernel density estimation (green for interoception, orange for exteroception).

### Comparison of the signal change for cardiac and gastric interoception in the insula

We directly compared the signal change in the insular subregions for cardiac and gastric interoception. Averaged parameter estimates were extracted from the insular ROIs independently for cardiac and gastric interoception (i.e., 12 values for each condition). We then conducted a repeated-measures ANOVA, with anatomical location and condition as the within-factors, independently for each hemisphere. The results indicated that, in both hemispheres, the main effect of anatomical location and the interaction between location and condition were significant (left, *F*_5, 150_ = 56.37, *η*_*p*_ = .65, *F*_5, 150_ = 8.89, *η*_*p*_ = .22; right, *F*_5, 150_ = 39.18, *F*_5, 150_ = 15.42, *η*_*p*_ = .34; all *p*s < .001), but the main effect of condition was not (left, *F*_1, 30_ = 0.07, *η*_*p*_ = .00; right, *F*_1, 30_ = 0.15, *η*_*p*_ = .01; *p*s > .70) (Figure 4C), replicating the results of previous study (Haruki and Ogawa, 2021). Post-hoc analysis of the left ROIs revealed that the ALG (corresponding to the dorsal posterior insula) showed a marginally significant activation higher in gastric than cardiac interoception (F_1, 30_ = 3.83, corrected *p* = .059). Post-hoc analysis of the right ROIs also revealed that the right ASG and MSG exhibited higher activation in cardiac than gastric interoception (F_1, 30_ = 5.93, corrected *p* = .021; F_1, 30_ = 6.57, corrected *p* = .016, respectively) while marginally significant activation higher in gastric interoception was found in the ALG (F_1, 30_ = 4.13, corrected *p* = .051).

Furthermore, we found that a differences in averaged activation between interoception and exteroception; the ANOVA on activations in the left insula revealed that significant effects of anatomical location (F_5, 150_ = 52.71, p < .001, *η*_*p*_ = .64) and interaction between condition and location (F_5, 150_ = 13.45, p < .001, *η*_*p*_ = .31) (Extended Data Figure 4-1). Post-hoc analysis showed that the left MSG and PSG were more activated by interoceptive attention (F_1, 30_ = 7.38, p = .011, *η*_*p*_ = 20; F_1, 30_ = 18.70, p < .001, *η*_*p*_ = .38, respectively). The ANOVA on the right insula activation revealed significant effects of anatomical location (F_5, 150_ = 41.02, p < .001, *η*_*p*_ = .58) and interaction between condition and location (F_5, 150_ = 12.23, p < .001, *η*_*p*_ = .28) as well. We found that exteroceptive attention more activated the right ASG than interoceptive attention by post-hoc analysis (F_1, 30_ = 10.28, p = .003, *η*_*p*_ = .26).

## Discussion

Our results empirically demonstrate that cardiac and gastric interoception have similar but distinct neural substrates. Direct comparison of brain activation revealed that cardiac and gastric interoception activated similar brain regions including the insula, frontal operculum, parietal operculum, middle cingulate, and supplementary motor area, when contrasted to the control visual condition. However, compared to gastric interoception, cardiac interoception activated the right anterior insula extending to the frontal operculum more; gastric interoception enhanced activations in the occipitotemporal visual cortices, bilateral primary motor, primary somatosensory, left orbitofrontal, and bilateral posterior hippocampus. Moreover, our MVPA-based ROI analyses revealed that the left middle-anterior insula had a distinct neural representation for cardiac and gastric interoception.

Previously, heartbeat perception has been studied as a representative of interoception in general, suggesting the right anterior insula as the most relevant region for generating subjective experience of internal bodily states (Craig, 2009; Critchley and Garfinkel, 2017). We, however, found that cardiac interoception activated the right anterior insula more than gastric interoception, suggesting that the right anterior insula may dominantly code cardiac interoception, rather than the interoception arising from other sources (e.g., gastric). In line with this idea, the most consistent activation in the right insula has been found by a meta-analysis focusing on cardiac interoception (Schulz, 2016). Furthermore, previous studies have found a robust correlation between individual accuracy of cardiac interoception and heightened activation of the right anterior insula (Caseras et al., 2013; Critchley et al., 2004; Haruki and Ogawa, 2021; Pollatos et al., 2007), but this was not the case for awareness of breathing (Wang et al., 2019) or skin conductance (Baltazar et al., 2021). The activation dominance in the right anterior insula may be explained by its functional roles: people can notice increased arousal with cardiac interoception (Paulus et al., 2019). The right insula appears essential to perceive arousal as the resection of the region causally diminishes physiological and emotional arousal (Holtmann et al., 2022; Terasawa et al., 2021). Moreover, presenting accelerated cardiac feedback increases perceived physiological arousal (Story and Craske, 2008) and, more importantly, activates the right anterior insula (Gray et al., 2007; Kleint et al., 2015). These findings, in conjunction with the present results, imply that cardiac interoception closely involves physiological arousal and thus activates the right anterior insula by merely focusing on cardiac sensation. To sum up, the relationship between the right anterior insula and interoception appears stronger in cardiac interoception than in other modalities; such dominance of cardiac interoception may reflect the functional role of cardiac interoception that informs physiological arousal.

We also found that the brain regions underlying gastric interoception encompassed the visual cortex compared to cardiac interoception. The involvement of the visual cortices in gastric interoception appears puzzling, but converging evidence suggests a strong connection between the visual cortex and stomach function. Recently, Rebollo and colleagues found temporally delayed connectivity between brain activity and the intrinsic electrical rhythm generated by the stomach, which includes the occipitotemporal visual cortices in addition to the somatosensory and motor areas (Rebollo et al., 2018; Rebollo and Tallon-Baudry, 2022). Crucially, the visual areas activated in the current experiment largely overlapped with the clusters included in the gastric network. Other past research shows that electrical or vibration stimuli on the stomach evoked neural activation in the visual area of the occipital area in rats (Cao et al., 2019), cats (Pigarev et al., 2013), and even humans (Mayeli et al., 2021). Considering all these reports, gastric functions are strongly tied to the visual cortex; here, we demonstrated that gastric interoception activated the visual cortex even without any direct stimulation of the stomach.

The reason for the involvement of the visual area may be that gastric interoception is closely related to foraging and feeding behavior that requires the integration of visuospatial information with energy status and motor coordination (Kanoski and Grill, 2017; Rebollo et al., 2018). Presumably, hungry people would change their sensory sampling and behavior in the external world to maximize the chance of food consumption. The current results support this idea; we observed activation related to gastric interoception in the left orbitofrontal cortex, bilateral hippocampus, primary motor cortex, and visual areas, which are all associated with food intake. For example, the orbitofrontal cortex has been suggested to modulate eating behavior by encoding the nutritional and reward values for food or food cues (Seabrook and Borgland, 2020). A meta-analysis of fMRI studies of viewing food pictures compared with non-food pictures found the left orbitofrontal cortex as the most consistent activation for visual food stimuli (van der Laan et al., 2011). Furthermore, the role of the hippocampus in controlling food intake and regulating energy status has received considerable attention in the past few years (Quigley et al., 2021; Suarez et al., 2019); recent evidence indicates that the human hippocampus encodes ongoing nutritional states in response to food cues (Jones et al., 2021). Based on these findings, we consider that neural responses to gastric interoception could be encoded in relation to food intake, which was activated by focusing on bodily sensations without any stimulation or food-related cues in the present study.

The present ROI analyses revealed that the insula had distinct representations for cardiac and gastric interoception. Only the right ASG and MSG, corresponding to the dorsal anterior insula, showed higher activation in cardiac than gastric interoception; in the right mid-posterior and left insula, there was no activation difference. Nevertheless, we identified the left PSG, corresponding to the dorsal middle insula, in different activation patterns for cardiac and gastric interoception. These results elaborate on how the human insula represents interoception; in particular, we suggest the left middle insula as coding the viscera-specific interoception for the first time. Interestingly, the posterior insula did not show separable activation for cardiac and gastric interoception nor higher activation than baseline. Researchers have considered the posterior insula as “primary” interoceptive cortex as it receives an initial cortical input of visceral signals (Craig, 2002; Evrard, 2019). The current inactivity of the posterior insula may support its role that encodes an ongoing change of bodily signals (Craig et al., 2000; Meier et al., 2018), rather than a subjective awareness of bodily states. This is because, in the interoceptive attention task, participants must be aware of interoception without homeostatic perturbation. Together, our ROI analyses would support the gradual process of bodily signals in the insula along the posterior-anterior axis (Craig, 2009): the posterior codes physical changes of the signal while the subjective differences in interoception would be represented in the middle-anterior insula.

Unfortunately, we did not record any physiological data during the fMRI scanning, potentially limiting the current study. Simultaneous recording of electrocardiogram (ECG) and electrogastrogram would address the interaction between brain activation and physiological change evoked by interoceptive attention. But, we consider that the lack of physiological recording does not challenge the validity of the present study because past research found that cardiac parameters (heart rate and ECG amplitude) did not differ by directing attention between interoception and exteroception (Petzschner et al., 2019). On a related note, previous studies have suggested that external processes such as cutaneous sensations could affect interoception (particularly in the cardiac domain) (Khalsa et al., 2009). Although we asked participants to focus on their internal sensations, we could not completely rule out the possibility that participants felt the heartbeat sensation from their skins. However, in the first place, interoception may not arise solely from a single visceral sensation— as when heartbeat perception is felt as the contraction of blood vessels and beating of the heart, or hunger is mediated by both the physical contents of the stomach and a chemical signal. If so, researcher may benefit from isolating the neural substrates of cardiac and gastric interoception localized in the present study from a visceral topography to reveal the process of (visceral) multisensory integration. Another possible limitation is that the difference in task difficulty in each condition could affect the results. Although we tried to make the difficulty comparable among each condition, there was a significant effect of the task condition on the subjective intensity of stimulus—subjective intensity of the visual stimulus was rated the highest while that of the stomach sensation the lowest of the three. We thus performed another analysis that excluded the effect of the subjective rating of stimulus intensity, indicating that the brain activation was almost unaffected by individual differences in the subjective rating. The results support our original ideas that the present brain activation in cardiac and gastric interoception was elicited by the object (the heartbeat and stomach sensation) of attentional focus not the difference in task difficulty. Recording a trial-by-trial fluctuation of the subjective intensity of stimulus would strengthen the present findings that the subjective differences in interoception are encoded differently in the brain.

## Acknowledgement

YH is a research fellow of Japan Society for the Promotion of Science (JSPS) concurrently. We would like to thank Editage (www.editage.com) for English language editing.

## Notes

**Conflict of Interest** We declare no conflicts of interest associated with this paper.

**Funding sources** This work was supported by JSPS KAKENHI Grant Number 20K20423 to KO and 22J20351 to YH and Graduate Grant Program of Graduate School of Humanities and Human Sciences, Hokkaido University, to YH.

https://osf.io/8pg37/

### Competing Interest Statement

The authors have declared no competing interest.

### Summary of Updates

Sections on Introduction updated to strengthen the validity of our experimental paradigm; Sections on Methods and Results updated according to our reanalyses; Figure 2 updated to show overlapping brain regions more clearly; Figure 3 has been created to show brain activation of interoception contrasted to control condition; Figure 4-1 has been created to support Figure 4 (original Figure 3); Data that were used to create the Figures and Tables are now available.

